# Meta-analysis of genome-wide DNA methylation and integrative OMICs in human skeletal muscle

**DOI:** 10.1101/2020.09.28.315838

**Authors:** S Voisin, M Jacques, S Landen, NR Harvey, LM Haupt, LR Griffiths, S Gancheva, M Ouni, M Jähnert, KJ Ashton, VG Coffey, JM Thompson, TM Doering, A Gabory, C Junien, R Caiazzo, H Verkindt, V Raverdy, F Pattou, P Froguel, JM Craig, S Blocquiaux, M Thomis, AP Sharples, A Schürmann, M Roden, S Horvath, N Eynon

## Abstract

Knowledge of age-related DNA methylation changes in skeletal muscle is limited, yet this tissue is severely affected by aging in humans. Using a large-scale epigenome-wide association study (EWAS) meta-analysis of age in human skeletal muscle from 10 studies (total n = 908 human muscle methylomes), we identified 9,986 differentially methylated regions at a stringent false discovery rate < 0.005, spanning 8,748 unique genes, many of which related to skeletal muscle structure and development. We then integrated the DNA methylation results with known transcriptomic and proteomic age-related changes in skeletal muscle, and found that even though most differentially methylated genes are not altered at the mRNA or protein level, they are nonetheless strongly enriched for genes showing age-related differential expression. We provide here the most comprehensive picture of DNA methylation aging in human skeletal muscle, and have made our results available as an open-access, user-friendly, web-based tool called *MetaMeth* (https://sarah-voisin.shinyapps.io/MetaMeth/).

## Introduction

While human lifespan (i.e. the number of years alive) has increased by ~3.5 years/decade since 1900^1^, healthspan (i.e. number of years spent in good health) has not increased to the same extent. In 2015, people lived 5 years longer than in 2000, but only 4.6 years longer in good health^2^. Ageing leads to the progressive loss of muscle mass and strength, resulting in a disorder termed sarcopenia. Sarcopenia is a serious condition leading to an increased risk of many conditions including cancer, type-2 diabetes, and cardiovascular diseases^3^. This process is driven by a host of adverse molecular changes in skeletal muscle with advancing age. Unravelling the molecular changes caused by aging in skeletal muscle is the basic foundation for the development of drugs and targeted health-related interventions to help prevent sarcopenia, and maximise healthspan.

Epigenetics are modifications of DNA that confer on the cell the ability to remember a past event^4^. Epigenetic changes are one of the *primary* hallmarks of aging, leading to dysregulated nutrient sensing, mitochondrial dysfunction and cellular senescence, which ultimately results in stem cell exhaustion and altered intercellular communication^5^. The best characterized epigenetic modification in the context of aging is DNA methylation. DNA methylation occurs at millions of CpG dinucleotides in the genome and changes considerably with age in various human tissues^6^, including skeletal muscle^7–9^. Age-related DNA methylation changes in skeletal muscle may be one of the molecular mechanisms underlying sarcopenia, but the full picture is fragmentary. To date, four epigenome-wide association studies (EWAS)^7,8,10,11^ have probed age-related DNA methylation changes in the muscle methylome, and both relied on relatively small sample sizes (n=10-50). Studies relying on a small sample size fail to detect small effect sizes and can be prone to large error, so larger initiatives are needed to identify the comprehensive list of CpG loci that change in methylation with age in human skeletal muscle. Meta-analysis, significantly increase statistical power and are more likely to identify robust age-related methylation sites^12^. Current understanding of epigenetic aging in skeletal muscle also remains incomplete as insight into the functional consequences of age-related epigenetic changes remain poorly understood. Whether age-related changes in DNA methylation in muscle cause or stem from changes in mRNA and protein expression is currently unknown.

To address these gaps, we performed a large-scale bioinformatics analysis of DNA methylation, mRNA and protein changes with age in human skeletal muscle. We integrated original DNA methylation data from our laboratory (the Gene SMART cohort) with available open-access data from multiple repositories and published studies. Firstly, we aimed to identify robust age-related CpGs in skeletal muscle in an EWAS meta-analysis of age, combining n = 908 samples from 10 datasets. Second, we performed enrichment analyses to unravel the potential functional consequences of these robust age-related DNA methylation changes. Thirdly, we integrated age-related methylome changes with transcriptome and proteome changes in skeletal muscle using two external, large-scale studies. Finally, we updated our skeletal muscle epigenetic clock ^9^ with an additional 371 samples, reaching a total of 1,053 human skeletal muscle methylomes from 16 datasets. Importantly, we have made the results of our analysis available as an open-access, user-friendly, interactive web-based tool, *MetaMeth* (https://sarah-voisin.shinyapps.io/MetaMeth/), enabling users to look at age-related changes in any gene of interest across the muscle methylome, transcriptome and proteome.

## Results

### Widespread age-related DNA methylation changes at genes involved in skeletal muscle structure, development and function

We first conducted a powerful EWAS meta-analysis of age in skeletal muscle using 10 datasets (total n = 908 samples), and uncovered a small, widespread effect of aging on the skeletal muscle epigenome. Eight percent of all tested CpGs were associated with age in skeletal muscle (61,006 differentially methylated positions (DMPs) corresponding to 9,986 differentially methylated regions (DMRs), both at FDR < 0.005, **Fig 1A** and Supplementary Tables 2 & 3). We found even numbers of hypomethylated and hypermethylated DMPs (50% hypo- and 50% hyper-DMPs, Supplementary Table 2) and the magnitude of age-related DNA methylation changes was small and similar for both hypo- and hyper-DMPs: hypo-DMPs lost an average of ~7% in methylation per decade of life, and hyper-DMPs gained an average of ~6% in methylation per decade of life (**Fig 1B**).

**Figure 1.**
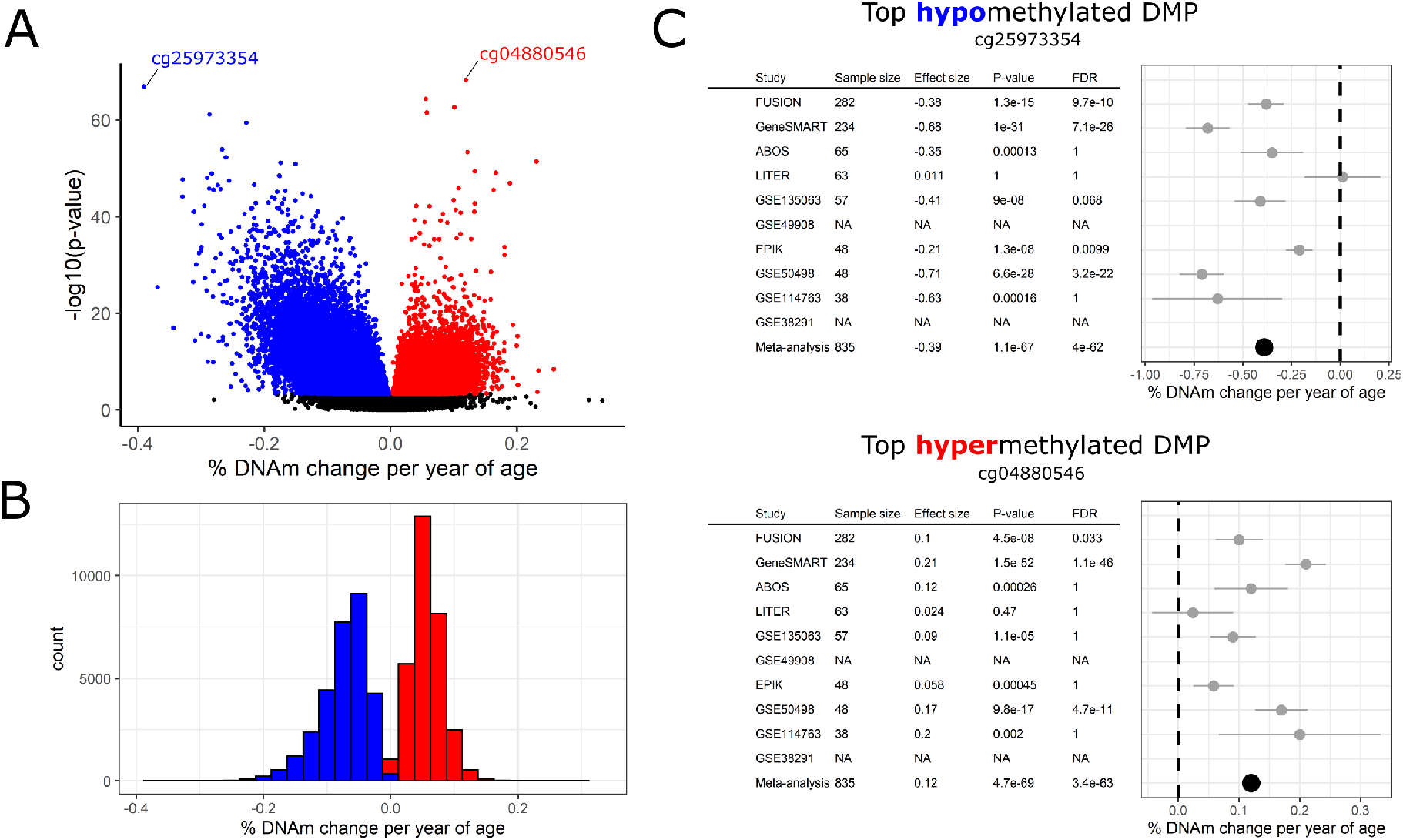
Age-related DNA methylation loci in human skeletal muscle. A) Meta-analysis effect size (x-axis) and meta-analysis significance (y-axis) for the 723,655 tested CpGs. Hypomethylated (blue) and hypermethylated (red) points represent differentially methylated position (DMPs) at a false discovery rate (FDR) < 0.005. B) Distribution of age-related DNA methylation change at hypo- and hyper-DMPs. C) Forest plots of the top hypo- and hyper-methylated DMPs, showing sample size, effect size, p-value and FDR for each individual study as well as their meta-analysis. Studies with missing information (“NA”) mean that this CpG was not analysed in the dataset.

Each dataset had a unique study design that required adjustment for factors that are known to affect DNA methylation levels, such as sex^13^, body mass index (BMI)^14^, and type 2 diabetes (T2D)^15^. We adjusted each dataset for these factors, but we noted that age was associated with BMI or T2D in some datasets (Supplementary Table 1). For example, older individuals from the GSE50498 dataset had a BMI than younger individuals (4.1 kg/m^2^ heavier, p = 0.0011) so it is possible that the age-related signal captured in this dataset was partially confounded by BMI. We repeated the meta-analysis without GSE50498, but results were largely unchanged (Supplementary Fig 3A). We also repeated the meta-analysis excluding T2D patients from the FUSION, ABOS and GSE38291 datasets, but results remained unchanged (Supplementary Fig 3B). Finally, we repeated the meta-analysis without the ABOS dataset whose muscle of origin differed from that of the other datasets (*rectus abdominis* vs *vastus lateralis* muscle). However, results remained unchanged (Supplementary Fig 3C). This confirms that our results are not confounded by BMI, T2D or the type of skeletal muscle.

We then focused on the DMRs for all downstream analyses, as DMRs remove spatial redundancy (CpG sites within ~500 bp are typically highly correlated^16^), and they may provide more robust and functionally important information than DMPs^17,18^. As with DMPs, we found even numbers of hypomethylated and hypermethylated DMRs (52% hypo- and 48% hyper-DMRs, Supplementary Table 3). DMRs’ distribution in chromatin states was different from that of all tested CpGs (χ^2^-test p-value < 2.2 × 10^−16^, **Fig 2**). DMRs were strongly under-represented in quiescent regions, while over-represented at enhancers and around active transcription start sites (TSS). However, hypo-DMRs were more strongly over-represented in genic enhancers and around active TSS; conversely, only hyper-DMRs showed overrepresentation in and around bivalent enhancers and promoters, and in regions actively repressed by PolyComb proteins. The distribution of hyper- and hypo-DMRs also varied with respect to CpG islands: both were under-represented in open seas and over-represented in CpGs island shores, but only hyper-DMRs were over-represented in CpG islands (χ^2^-test p-value < 2.2 x 10^−16^, **Fig 2**). Finally, both hypo- and hyper-DMRs were under-represented in CCCTC-binding factor (CTCF) binding sites in differentiated skeletal muscle myotubes, but only hyper-DMRs were strongly over-represented in enhancer of zeste homolog 2 (EZH2) binding sites (**Fig 2**).

**Figure 2.**
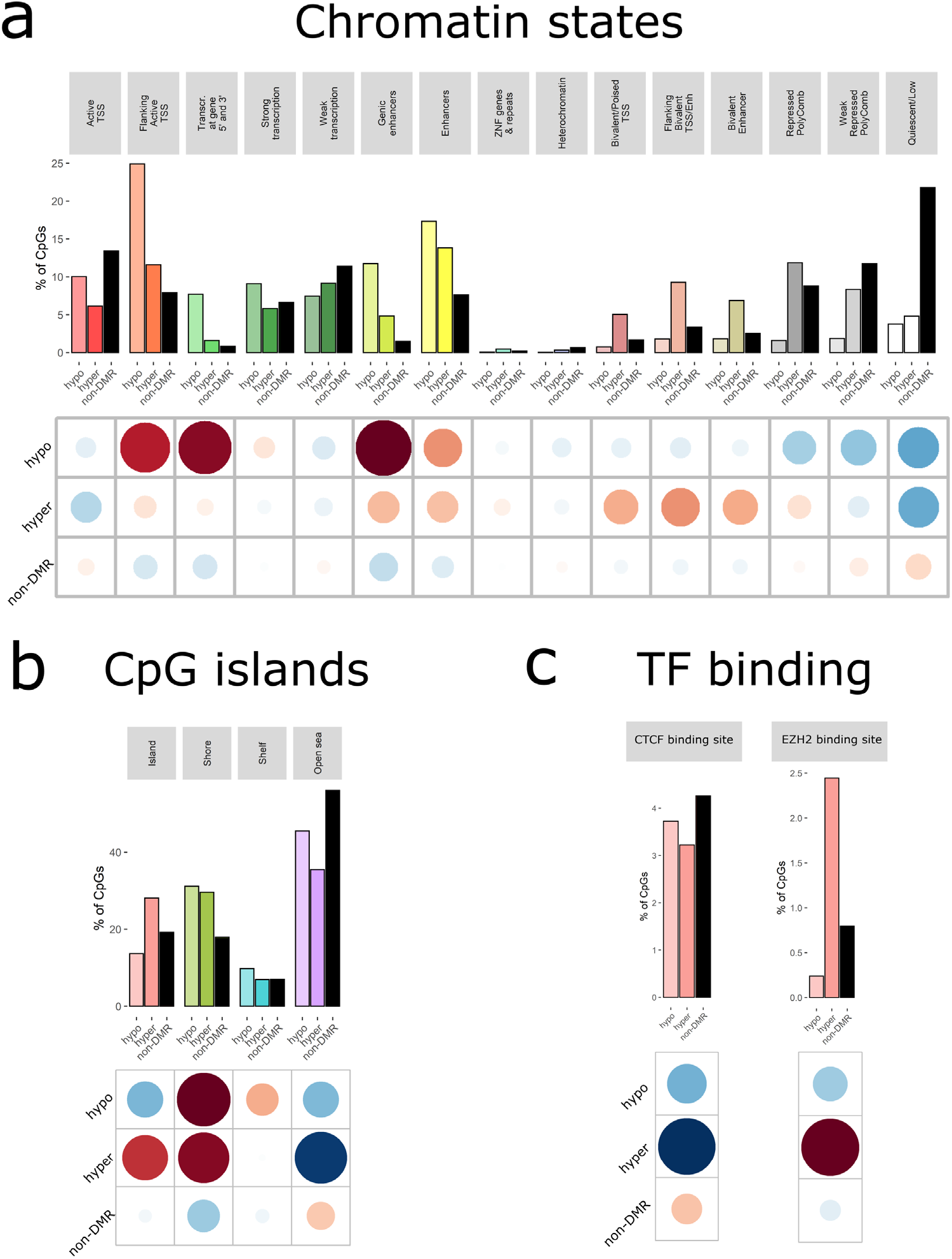
Distribution of hypomethylated, hypermethylated differentially methylated regions (DMRs) and non-DMRs in functional regions of the genome. Distribution in chromatin states from male skeletal muscle from the Roadmap Epigenomics Project^19^ (a); distribution with respect to CpG islands (b), shore = ± 2kb from the CpG island, shelf = ± 2-4 kb from the CpG island, open sea = > 4kb from a CpG island; distribution in CCCTC-binding factor (CTCF) and enhancer of zeste homolog 2 (EZH2) binding sites in skeletal muscle myotubes differentiated from the HSMM cell line (HSMMtube) from the ENCODE project (c). The grids under the figures represent the residuals from the χ2-test, with the size of the circles being proportional to the cell’s contribution; red indicates an enrichment of the DMR category in the functional region, while blue indicates a depletion of the DMR category in the functional region.

Next, we integrated a comprehensive annotation of Illumina HumanMethylation arrays^20^ with chromatin states from the Roadmap Epigenomics Project^19^ and the latest GeneHancer information^21^ to map the DMRs to genes (Supplementary Table 3). Including non-coding genes, there were 8,748 genes that harboured at least one DMR, hereinafter referred to as differentially methylated genes (DMGs). A pathway enrichment on the DMRs revealed that DMGs were enriched for 158 GO terms (Supplementary Table 4), many of which related to skeletal muscle structure development, muscle contraction, and calcium transporter regulation (**Fig 3**). However, we found no enrichment for any KEGG term (all FDR > 0.005).

**Figure 3.**
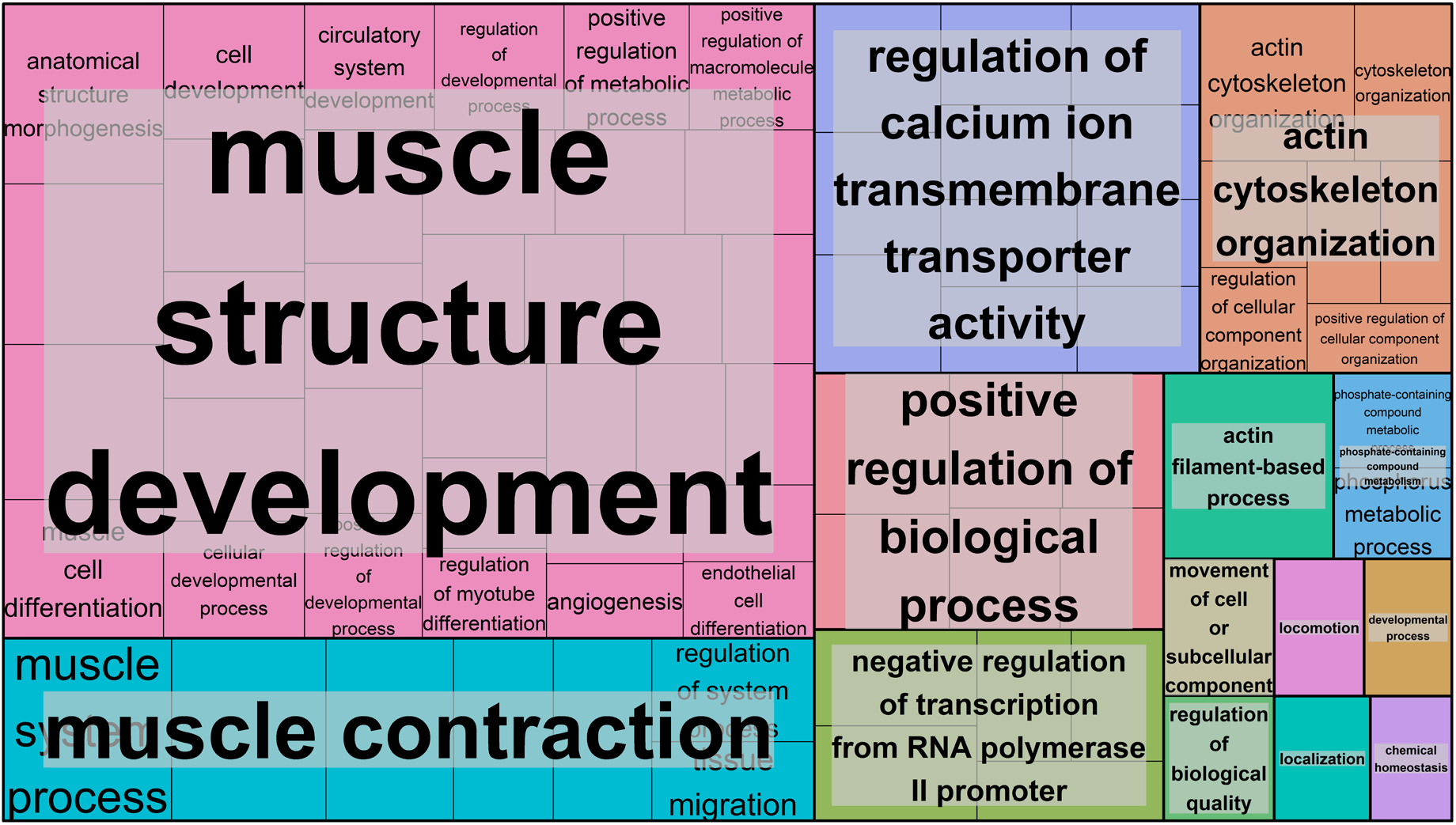
Gene Set Enrichment Analysis of the differentially methylated genes. This treemap shows the clustering of the 131 significant Gene Ontology (GO) terms belonging to the “biological processes” category. The 131 GO terms were clustered based on semantic similarity measures using REVIGO^22^, with each rectangle corresponding to a single cluster representative. The representatives are joined into ‘superclusters’ of loosely related terms, visualized with different colours. The size of the rectangles is proportional to the –log^10^(p-value) of the GO term.

### Differentially methylated genes are enriched for genes showing age-related changes at the mRNA and protein levels

We investigated the potential downstream effects of these age-related DNA methylation changes on mRNA and protein expression in muscle. We utilised two external published studies: a transcriptomic meta-analysis of age that combined 2,852 public gene expression arrays in skeletal muscle^23^, and a large-scale proteomic analysis of age in skeletal muscle from 58 healthy individuals aged 20 to 87 years^24^. Su *et al*.^23^ identified 957 genes whose mRNA levels change with age, and Ubaida-Mohien *et al.*^24^ identified 1,265 genes whose protein levels change with age. Fourty-one percent of the genes whose mRNA levels change with age were also altered at the DNA methylation level, and 42% of the genes whose protein levels change with age were also altered at the DNA methylation level (**Fig 4A**). Furthermore, the DMGs included proportionally many more differentially expressed genes than the non-DMGs (χ^2^-test p-value < 2.2 × 10^−16^, **Fig 4A**), indicating that such a large overlap between differential DNA methylation and differential gene expression with age cannot be attributed to chance alone.

**Figure 4.**
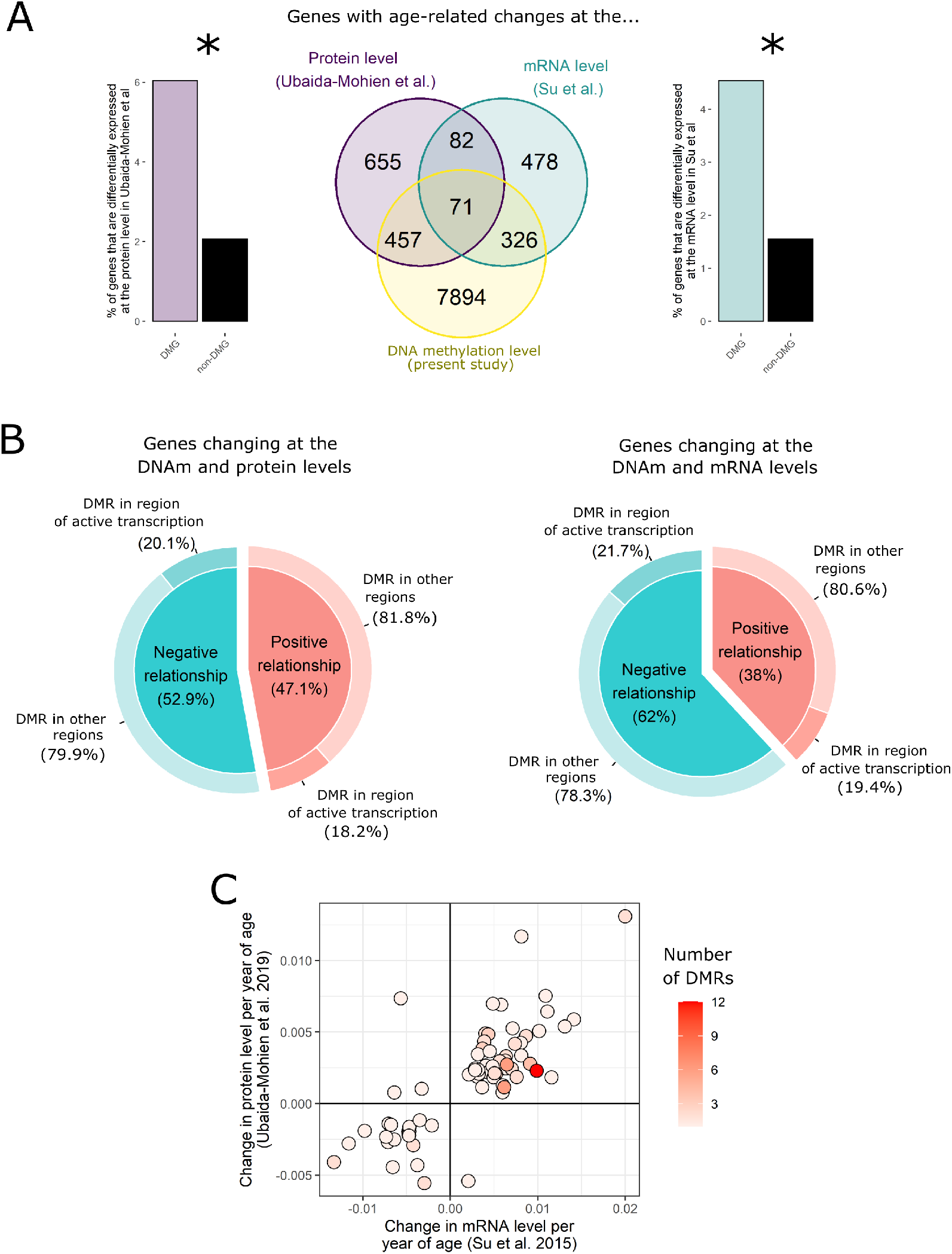
Integration of DNA methylation, mRNA, and protein changes with age in human skeletal muscle. A) Overlap between genes that change with age at the DNA methylation level (yellow, present study), mRNA level (green, Su et al. 2015^23^) and protein level (purple, Ubaida-Mohien et al. 2019^24^). On each side of the Venn diagram, we showed the distribution of differentially expressed genes among the differentially methylated genes (DMGs) and the non-differentially methylated genes (non-DMGs). *χ^2^-test p-value < 0.005. B) Relationship between age-related DNA methylation changes and mRNA changes (right) or protein changes (left): “negative relationship” means that a gene that was upregulated with age at the gene expression level showed lower DNA methylation with age in the present study, and a gene that was downregulated with age at the gene expression level showed higher DNA methylation with age in the present study. As the relationship between DNA methylation and gene expression differs depending on the genomic context, we further split the age-related DNA methylation changes between those located in regions of active transcription and those located in other regions. C) Scatterplot showing the change in mRNA (x-axis) and protein (y-axis) per year of age for the 71 genes altered at all three omics levels. Each gene was coloured according to the number of DMRs annotated to it, from 1-3 DMRs for most genes all the way up to 12 DMRs. Naturally, longer genes (e.g. NXN, ABLIM2) have a greater propensity to have more DMRs given their high numbers of CpGs.

Next, we investigated in more details the relationship between DNA methylation and mRNA or protein expression. This relationship is complex and depends on the genomic context, particularly the underlying chromatin state^25^; an increase in DNA methylation is usually associated with a downregulation of gene expression, but the opposite pattern is observed in gene bodies of actively transcribed genes. We found that the relationship between DNA methylation and mRNA expression was mostly negative, regardless of whether the DMR was in a gene body or not, but the relationship between DNA methylation and protein expression did not show any predominant pattern (**Fig 3B**).

Seventy-one genes were altered at all three omic levels (Supplementary Table 5, **Fig 3C**). Moreover, there was a high concordance between the transcriptomic and proteomic studies: an age-related increase in mRNA level was most often mirrored by an age-related increase in protein level, and vice versa (**Fig 3C**).

### *MetaMeth*: an online tool to visualise the aging profile of human skeletal muscle

We have made the results of the EWAS meta-analysis of age in skeletal muscle available as an online webtool called *MetaMeth* (https://sarah-voisin.shinyapps.io/MetaMeth/). The homepage of the website provides a detailed list of instructions on how to visualise results and focus on specific CpGs, genes, or genomic regions of interest in a user-friendly, interactive manner. To obtain forest plots for individual CpGs, users can enter the name of their CpG of interest (e.g. “cg11109027”) in the “Forest Plot” tab and the corresponding graph will appear, with the possibility to download the plot in jpg, png or tif formats and at any resolution. To help with choosing CpGs to display, users can filter the list of CpGs based on their genomic location (e.g. genomic region, annotated gene, position with respect to CpG islands, chromatin states in male and female skeletal muscle, and TF binding). To download summary statistics for DMPs or DMRs in a table format, users can go to the “Summary Tables” tab, and download the data as an excel or csv file, after optionally filtering data based on genomic location and statistics. Finally, we have also displayed the scatterplot of genes showing methylation, mRNA and protein changes with age as an interactive graph: users simply need to hover their mouse on one point of the graph to be shown the name of the gene and the number of DMRs annotated to it. The code used to produce the website is available in open-access on Sarah Voisin’s Github account (https://github.com/sarah-voisin/MetaMeth).

### More samples in the muscle epigenetic clock does not change age prediction accuracy

The present EWAS meta-analysis of age utilised all of the datasets included in the original muscle epigenetic clock (MEAT) that we recently published, with the exception of datasets that were invariant in age and the datasets that were too small (n < 20) (see Material & Methods)^9^. The present study included an additional 371 samples from five datasets. Using the same algorithm and methodology, we updated the MEAT clock with these new samples, reaching a total of n = 1,053 human skeletal muscle samples from 16 datasets. The new version of the MEAT clock (MEAT 2.0) uses DNA methylation at 205 CpGs to predict age, 98 of which were in common with the first version of MEAT. We found that MEAT 2.0 only slightly outperforms the original version of MEAT, with an average Pearson correlation coefficient of 0.67 across datasets (vs 0.62 for the previous version of MEAT^9^) and a median error of only 4.5 years across datasets (vs 4.6 years for the previous version of MEAT^9^) (**Fig 5**).

**Figure 5.**
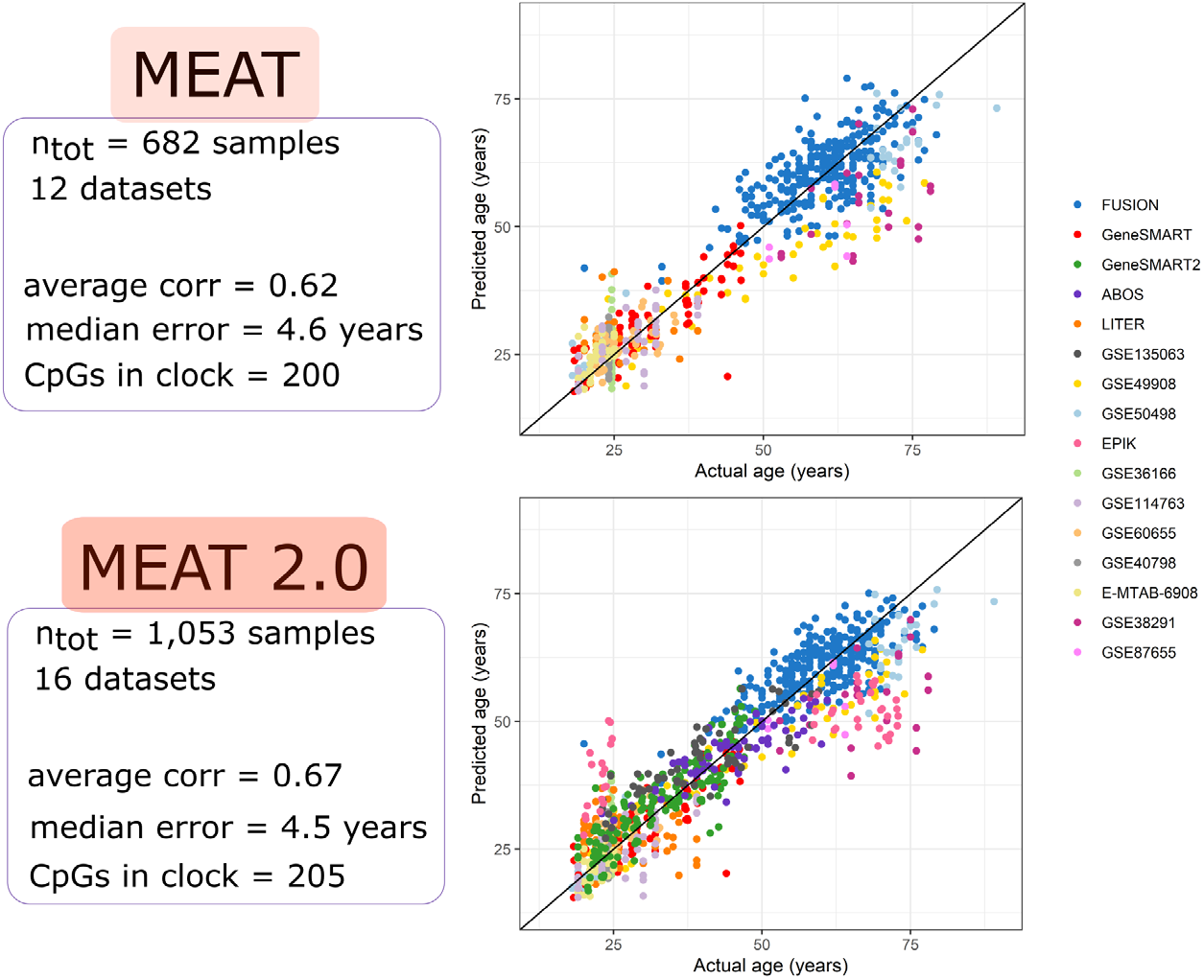
Original and new version of the muscle clock (MEAT). The top panel shows the original version of the muscle clock^9^ (MEAT) and the bottom panel the new version of the muscle clock (MEAT 2.0). The graphs on the right hand-side show predicted (y-axis) against actual (x-axis) age for each sample in the 16 datasets used to build the clocks. A leave-one-dataset out cross-validation (LOOCV) procedure was used to obtain predicted age for a given dataset in an unbiased manner (16 LOOCV were performed, one per dataset). The summary statistics reported on the left hand side are the average correlation between actual and predicted age across datasets, the median absolute error in age prediction across datasets, and the number of CpGs automatically selected by the algorithm to build the clock.

## Discussion

To paint a comprehensive picture of age-related DNA methylation changes in human skeletal muscle, we conducted a powerful EWAS meta-analysis of age in human muscle across the lifespan, combining 908 samples from 10 independent datasets. In this study, we were able to demonstrate a profound effect of age on the muscle methylome. Additionally, we have provided a detailed account of the genomic context of age-affected regions, reported putatively affected pathways, and integrated methylome changes with known transcriptome and proteome changes in muscle. To maximise the usefulness of this large-scale EWAS meta-analysis to the scientific community, we created a website named MetaMeth (https://sarah-voisin.shinyapps.io/MetaMeth/), which allows researchers to visualise results in an interactive and user-friendly manner. Finally, we updated our muscle clock^9^ with 371 newly acquired DNA methylation samples and found that the first version of the MEAT clock was already at optimal prediction accuracy.

Previous studies on the overall pattern of age-related DNA methylation changes in muscle showed mixed results, three reporting more hypermethylation with age^7,8,11^ and one finding slightly more hypomethylation with age^10^. We included three of these studies (GSE49908, GSE50498 and EPIK) in our meta-analysis, and found balanced amounts of hypo- and hyper-methylation. Differences in coverage between studies are unlikely to explain the discrepancy in results, since the three HumanMethylation arrays were represented in these studies (27k, 450k and 850k). It is more likely that the overall direction of age-related DNA methylation change became more nuanced once these small-scale studies were combined with the other nine datasets. This highlights the advantage of the meta-analysis approach we utilised in identifying robust ageing-related CpG sites across multiple, potentially conflicting studies. We detected thousands of age-related DMRs, likely thanks to the unprecedented power achieved with 908 human muscle samples. Age-affected regions were not randomly spread across the genome and were particularly abundant around active TSS regions and in enhancers. Furthermore, hypo- and hyper-methylated regions showed a distinct distribution largely consistent with previous reports on aging; during aging, DNA methylation tends to increase at Polycomb target genes^26,27^ and bivalent chromatin domains^27,28^, while decreasing at enhancers in both mice and humans^26,27^. To explain the age-related hypermethylation of Polycomb-target genes, Jung & Pfeifer proposed a mechanism involving competition between Polycomb complexes and DNA methyltransferases 3 (DNMT3)^29^: the ability of the Polycomb machinery to target unmethylated CpG-rich target sequences erodes with age, leaving room for DNMT3 to bind and slowly methylate Polycomb target genes over time, potentially leading to reduced plasticity of the hypermethylated genes. This was entirely consistent with our findings: hypermethylated DMRs were strongly enriched in CpG islands and EZH2 binding sites (EZH2 is the enzymatic subunit of the Polycomb complex). Polycomb-target genes and bivalent chromatin domains are linked to developmental and differentiation processes^26^, which corroborated the pathway enrichment showing numerous GO terms related to muscle cell differentiation and skeletal muscle development. Neither the root nor the functional consequences of enhancer hypomethylation are known, but it may stem from altered DNMT and TET enzymes activity, and might lead to activation of cryptic transcripts or disrupt enhancer-gene interactions^26^. Taken together, our findings indicate a widespread effect of age on DNA methylation levels in skeletal muscle at genes fundamental for skeletal muscle development, structure and differentiation.

We integrated the present EWAS meta-analysis of age with two large, published transcriptomic and proteomic studies of age in human skeletal muscle^23,24^. The differentially methylated genes showed an over-representation of genes known to change at the mRNA and protein level with age, pointing at a likely functional relationship between age-related DNA methylation and gene expression changes. Furthermore, genes whose mRNA levels increase with age were mostly hypomethylated in our meta-analysis, and vice-versa. We could not assess whether age-related DNA methylation changes are a cause or a consequence of age-related gene expression changes, but the two scenarios are not mutually exclusive. We also noted that age-related mRNA and protein changes in skeletal muscle were highly consistent, as there was a strong positive correlation between mRNA and protein changes with age in human skeletal muscle. This reinforces the utility of large-scale studies, including meta-analyses, to produce robust, replicable results identifying DNA methylation targets. Future studies should explore the origin and functional consequences of these age-related omic changes in human skeletal muscle, and investigate whether the cause of the aging processes is similar across tissues. As changes in the epigenetic landscape are one of the primary hallmarks of aging, understanding its origin would narrow down our focus on putative genetic or/and epigenetic regions, with the ultimate goal of targeting them with lifestyle or pharmacological interventions to slow down the ageing process at the molecular level. Future studies should aim to find interventions easily accessible to a wide range of people, such as exercise training or dietary interventions, to slow down, or perhaps even reverse, age-related epigenetic changes in skeletal muscle.

Recently, we established an epigenetic clock for human skeletal muscle, using 682 samples from 12 datasets^9^. Here we updated this clock (MEAT 2.0) by using 1,053 samples from 16 datasets, particularly adding more female and middle-aged individuals that were underrepresented in MEAT. MEAT 2.0 automatically selected 205 CpGs for age prediction, only 98 of which were in common with the CpGs selected by MEAT. While such a small overlap may seem surprising, it likely stems from the machine learning algorithm underlying the clocks: tens of thousands of CpGs change with age, but only a handful of CpGs are selected by the elastic net model, so this group of CpGs is only one of the many possible combinations of CpGs that can predict age with high accuracy^6^. We tested whether the accuracy of the muscle clock is improved by feeding more samples to the machine learning algorithm. Surprisingly, the accuracy of the new version of the clock barely improved, from 0.62 to 0.66 in average correlation between predicted and actual age, and from 4.6 to 4.5 years in median error in age prediction. This suggests that the original muscle clock was already sufficiently accurate for age prediction in human skeletal muscle using the Illumina HumanMethylation array technology. We have however updated the R package *MEAT* on Bioconductor with this new clock, providing users the possibility to choose between the old version (MEAT) and new version (MEAT 2.0) of the clock for their analyses.

The age-related changes in the muscle methylome uncovered herein, and the epigenetic age calculated from the MEAT clock reflect both intracellular changes in methylation levels, and age-related changes in muscle cell type composition. Older muscle tends to have a greater proportion of type I (slow-twitch) muscle fibres^30^, show fat^31^ and macrophage^32^ infiltration, and display lower numbers of satellite cells^33^, which can alter the methylome of bulk muscle tissue. However, we adjusted the analyses for bias and inflation^34^ to account for unmeasured factors such as population substructure, batch effects, and cellular heterogeneity^35^. Uncovering the intracellular changes of different muscle cell types with age was beyond the scope of this study, and we did not have information on individual cellular profiles to answer this question. Nevertheless, the results shown here, along with the epigenetic clock and open-access search engine we developed, may still be highly valuable to aging researchers whose focus is unrelated to cell-type-specific aging. It should also be noted that the conclusions of this study may not apply to the human population as a whole, as 98% of the samples were of Caucasian origin and 71% were from male subjects. Future studies should make efforts to profile the methylomes of underrepresented groups to provide a picture of aging that reflects the world population.

To provide the scientific community with a tool to assess DNA methylation changes with age in skeletal muscle, we have created a user-friendly, interactive, and transparent way to explore our results. We built a web-based tool called *MetaMeth* (https://sarah-voisin.shinyapps.io/MetaMeth/), largely inspired by the *MetaMex* tool developed by Pillon *et al*. for transcriptomic meta-analysis of exercise training and inactivity in human skeletal muscle^36^. Users are able to explore DMPs, DMRs, forest plots, and omics integration and to filter and download the results. This freely available website is likely to advance the field of aging science as a whole.

## Methods

### EWAS meta-analysis of age in skeletal muscle

We combined four datasets of genome-wide DNA methylation in skeletal muscle (the Gene Skeletal Muscle Adaptive Response to Training (SMART)^37^, the Limb Immobilisation and Transcriptional/Epigenetic Responses (LITER) study^9^, the Biological Atlas of Severe Obesity (ABOS) study^38^, and the Epigenetica & Kracht (EPIK) study^10^), with 5 datasets from the open-access Gene Expression Omnibus (GEO) platform (GSE49908^8^, GSE50498^7^, GSE114763^39^, GSE38291^40^, and GSE135063^41^), and the Finland-United States Investigation of NIDDM Genetics (FUSION) Study^42^ (phs000867.v1.p1). These summed up to a total of n = 908 skeletal muscle samples collected from men and women across the lifespan (age range 18-89 years old, Supplementary Fig 1, Supplementary Table 1). Samples were 98% Caucasian and 71% male (Supplementary Table 1). We excluded cohorts from our recently published paper^9^ with a narrow age range (age standard deviation < 5 years) as age-related differences in DNA methylation cannot be detected if age is invariant; we also excluded datasets with a limited number of samples (n < 20) for robustness. Samples from the Gene SMART cohort (n = 234) include two batches, and our recently published paper^9^ only includes the first batch of 75 samples available on the Gene Expression Omnibus platform (GSE151407). The additional 159 samples from the second batch include both men and women, before and after exercise intervention.

Details on the preprocessing of each DNA methylation dataset can be found elsewhere^9^. We conducted independent EWAS of age in skeletal muscle in each dataset, using robust linear models and moderated Bayesian statistics as implemented in *limma*^43^; to isolate the contribution of age to DNA methylation variability, we regressed DNA methylation level against age and adjusted, when the dataset included these covariates, for sex, BMI, diabetes status, batch, and timepoint (baseline/post intervention or training); we also added, when the dataset included repeated measures on the same individuals or related individuals, a random intercept using the duplicateCorrelation function to account for repeated measures from the same individuals or to account for twinship. We adjusted each EWAS for bias and inflation using the empirical null distribution as implemented in *bacon* (Supplementary Figure 2)^34^. Inflation and bias in EWAS are caused by unmeasured technical and biological confounding, such as population substructure, batch effects, and cellular heterogeneity^35^. The inflation factor is higher when the expected number of true associations is high (as it is for age); it is also greater for studies with higher statistical power^34^. The figures we found (Supplementary Fig 2) were consistent with the inflation factors and biases reported in an EWAS of age in blood^34^.

Results from the independent EWAS were combined using an inverse variance weighted meta-analysis with METAL^12^. We used METAL since it does not require all DNA methylation datasets to include every CpG site on the HumanMethylation arrays. Different sets of CpGs may be filtered out during preprocessing of each individual dataset, which means the overlap between the datasets is imperfect and a given CpG may only be present in 5 out of 10 datasets, or 8 out of 10 datasets. For robustness, we only included CpGs present in at least 6 of the 10 cohorts (723,655 CpGs). We used a fixed effects (as opposed to random effects) meta-analysis, assuming one true effect size of age on DNA methylation, which is shared by all the included studies. Nevertheless, Cochran’s Q-test for heterogeneity was performed to test whether effect sizes were homogeneous between studies (a heterogeneity index (I^2^) > 50% reflects heterogeneity between studies). The CpGs associated with age at a stringent meta-analysis FDR < 0.005 were considered DMPs. We then identified DMRs (i.e. clusters of DMPs with consistent DNA methylation change with age) using the *dmrcate* package, at a Fisher’s multiple comparison statistic < 0.005, a Stouffer score < 0.005 and a harmonic mean of the individual component FDRs < 0.005^44^. *dmrcate* works by smoothing the test statistic of CpGs separated by a maximum of 1,000 bp using a Gaussian kernel; then, it models the smoothed test statistics, computes and corrects p-values, and finally aggregates adjacent CpGs that are significant and within 1,000 bp of each other. We focused on the DMRs for all downstream analyses, as DMRs remove spatial redundancy (CpG sites within ~500 bp are typically highly correlated^16^), and they may provide more robust and functionally important information than DMPs^17,18^.

### Enrichment of DMRs in functional regions of the genome

We used a χ^2^-test to compare the distribution of hyper- and hypo-methylated DMRs with that of non-DMRs 1) at different positions with respect to CpG islands, 2) in different skeletal muscle chromatin states from the Roadmap Epigenomics Project^19^, and 3) in CTCF and EZH2 transcription factors binding sites in HSMMtube from the ENCODE project. CTCF is a multifunctional protein involved in gene regulation and chromatin organisation^45^, while EZH2 is the functional enzymatic component of the Polycomb Repressive Complex 2 (PRC2) ^46^. A p-value < 0.005 was deemed significant.

We performed GO and KEGG enrichment on the age-related DMRs using all tested CpGs as the background (i.e. the 723,655 CpGs included in the meta-analysis), thanks to the goregion function from the *missMethyl* package^47^. We used our own improved annotation of the epigenome and largely based on Zhou *et al*.’s comprehensive annotation of Illumina HumanMethylation arrays^20^ as well as the chromatin states in skeletal muscle from the Roadmap Epigenomics Project^19^, and the latest GeneHancer information^21^. The goregion function accounts for the biased distribution of CpGs in genes^48^. All GO and KEGG terms with FDR < 0.005 were deemed significant^49,50^. To make sense of the many GO terms obtained as output, we used REVIGO^22^ that clusters GO terms according to semantic similarity.

### Integration of methylome, transcriptome and proteome changes with age

Each gene with at least one DMR annotated to it was considered a differentially methylated gene (DMG). To gain insights into the functional consequences of DNA methylation changes with age in skeletal muscle, we compared DMGs with known differentially expressed genes at the transcriptomic^23^ and the proteomic^24^ levels with advancing age. A transcriptomic meta-analysis in skeletal muscle was recently published^36^, but it focused on exercise-induced changes instead of age-related changes. Thus, we used the transcriptomic meta-analysis of age by Su *et al*. that combined 2,852 public gene expression arrays in skeletal muscle and identified 957 genes whose mRNA levels changed with age^23^. Ubaida-Mohien *et al*. performed a large-scale proteomics analysis of human skeletal muscle and identified 1,265 genes whose protein levels were altered with age ^24^. We used a χ^2^-test to see whether a disproportionate number of DMGs were also differentially expressed at the mRNA or protein level, and a p-value < 0.005 was deemed significant.

### Update of the muscle epigenetic clock (MEAT 2.0)

Since the development of the original muscle clock that used 682 samples from 12 datasets to predict age from DNA methylation data ^9^, we gathered an additional 371 samples from 5 datasets (+159 from Gene SMART, +65 from ABOS, +42 from LITER, +57 from GSE135063 and +48 from EPIK). We therefore updated the clock with these new samples, using the same algorithm and methodology^9^. Briefly, we first preprocessed each dataset separately (i.e. probe/sample filtering, adjustment of type I and type II probes and correction for batch effects); then, we reduced each dataset to all the CpGs that were common between them (18,666 CpGs). To obtain DNA methylation profiles that were comparable between datasets, we calibrated each dataset to GSE50498 using an adapted version of the BMIQ algorithm^9^. We then used elastic net regression on a transformed version of age to create the new muscle clock (MEAT 2.0)^9^. Finally, given the limited number of datasets and the biased age distribution in each dataset, we estimated the accuracy of the new muscle clock in an unbiased manner using a leave-one-dataset-out cross-validation procedure, as described in our original paper^9^.

---

**Supplementary Figure 1.**
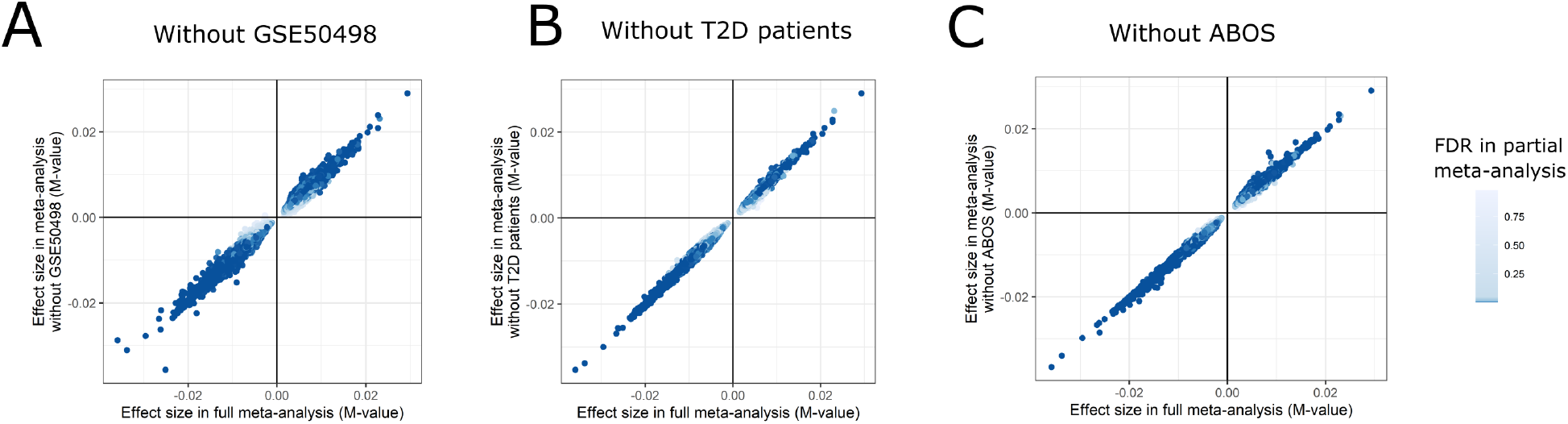
Comparison of results from the full meta-analysis and from a meta-analysis excluding GSE50498 (A), type 2 diabetes (T2D) patients (B) or the ABOS cohort (C). Each point is one of the 61,006 differentially methylated positions (DMPs) discovered in the full meta-analysis. To compare results from the full and partial meta-analyses, we plotted the effect size (change in M-value per year of age) in the full meta-analysis (x-axis), against the effect size in the partial meta-analysis (y-axis). To show whether DMPs remained significant in the partial meta-analysis, we colored points according to the false discovery rate (FDR) in the partial meta-analysis.

**Supplementary Figure 2.**
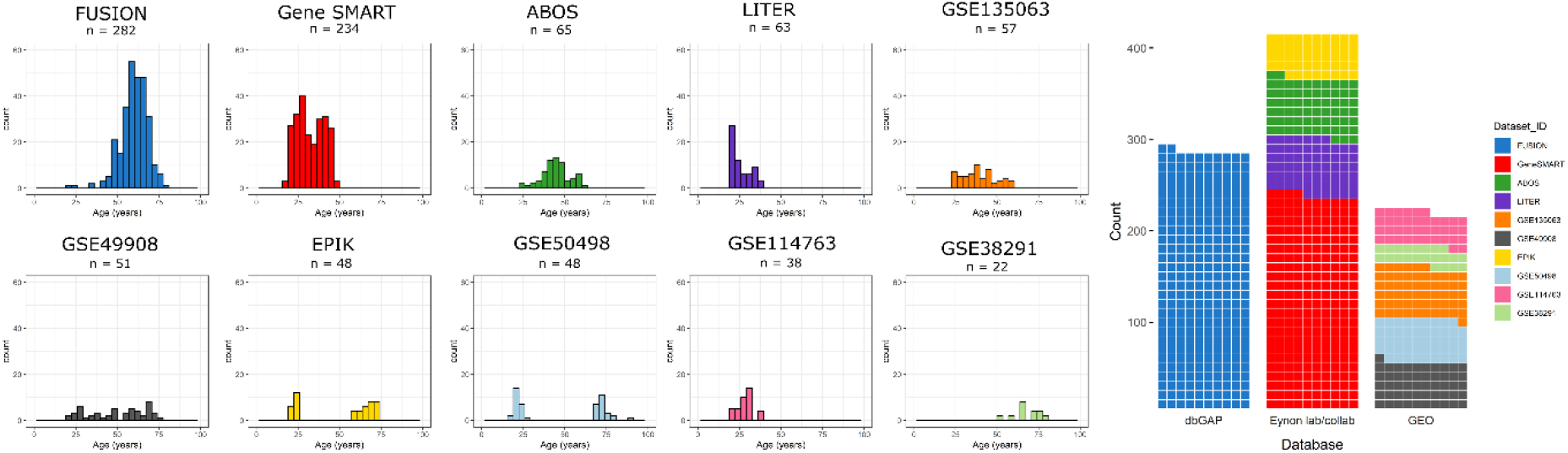
Age distribution in each of the 10 datasets included in the EWAS meta-analysis, and database of origin. dbGAP = database of Genotypes And Phenotypes; GEO = Gene Expression Omnibus

**Supplementary Figure 3.**
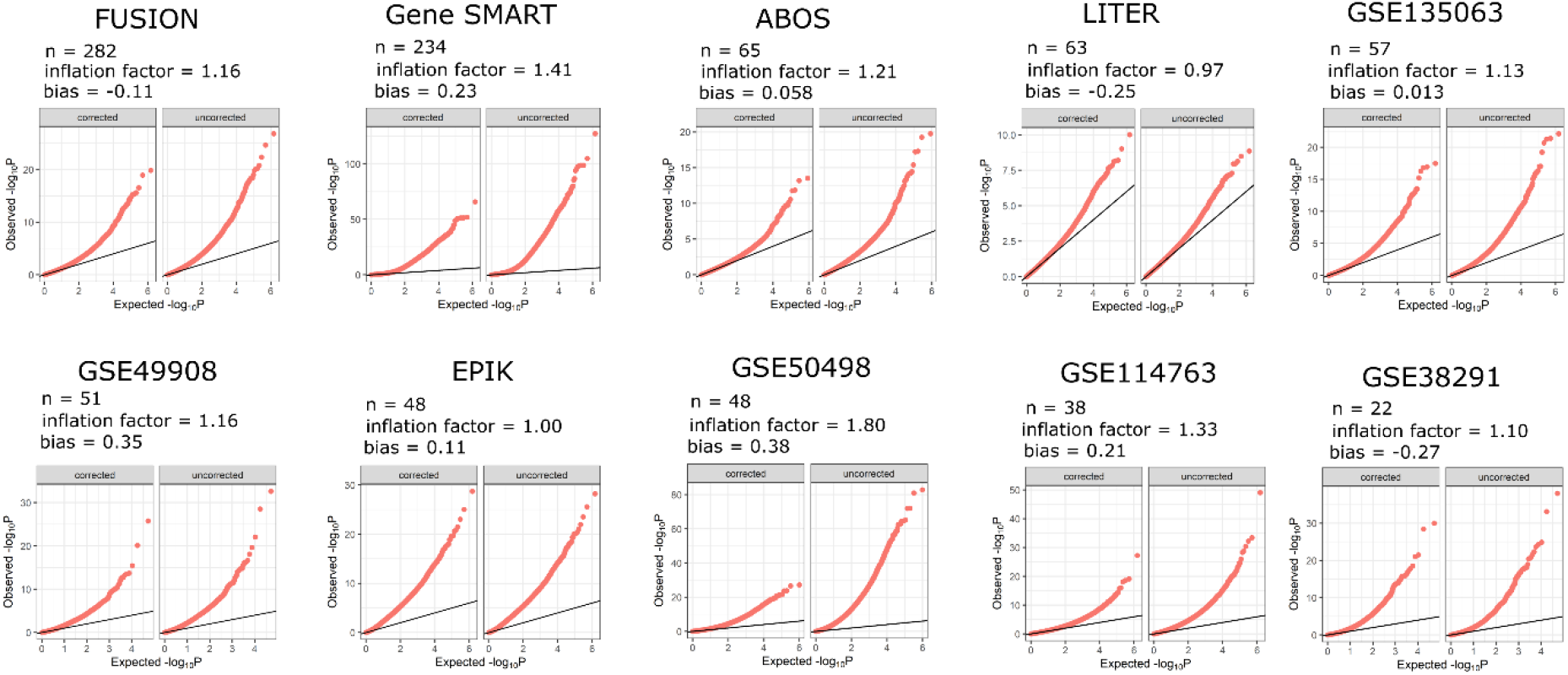
Quantile-quantile plot of −log^10^ transformed P-values for each of the 10 datasets included in the EWAS meta-analysis. Right panel using uncorrected P-values and left panel using bacon bias- and inflation-corrected P-values.

## Acknowledgements

We are grateful for the support of the Australian National Health & Medical Research Council (NHMRC) via Sarah Voisin’s Early Career Research Fellowship (APP11577321), and Nir Eynon’s Career Development Fellowship (APP1140644). We also thank the Australian Research Council (ARC) for supporting this study (DP190103081 and DP200101830), The Gene SMART and LITER studies were both supported by the Collaborative Research Network for Advancing Exercise and Sports Science (201202) from the Department of Education and Training, Australia. Mr Nicholas Harvey and Ms Jamie-Lee Thompson was supported by a PhD stipend also provided by Bond University CRN-AESS. This research was also supported by infrastructure purchased with Australian Government EIF Super Science Funds as part of the Therapeutic Innovation Australia - Queensland Node project (LRG). We also greatly acknowledge Erika Guzman at the ATGC/IHBI/QUT for performing the HMEPIC assays in the LITER study. Adam Sharples was supported by GlaxoSmithKline, North Staffordshire Medical Institute, The Society fort Endocrinology, the MRC and EPSRC, UK doctoral training centre, and the Norwegian School of Sports Sciences. The work was also supported by the German Ministry of Education and Research (BMBF: DZD grant 82DZD00302) and the Brandenburg State (Germany). The work was also supported by the German Ministry of Education and Research (BMBF: DZD grant 82DZD00302) and the Brandenburg State (Germany). The EPIK study was supported by the Foundation Scientific Research – Flanders (FWO Grant F.0898.15). We would also like to thank Mark Ziemann from Deakin University for his critical feedback on the manuscript.

## Contributions

Conceptualization: S.V., and N.E. Methodology: S.V., and S.H. Investigation: S.V. Formal analysis: S.V. Resources: M.J., S.L., N.R.H., L.M.H., L.R.G., S.G., M.O., M.J., K.J.A., J.M.T., A.G., C.J., R.C., H.V., V.R., F.P., P.F., S.B., M.T., A.P.S., A.S., M.R., S.H., and N.E. Software: S.V. Writing—Original draft: S.V., and N.E. Writing—Review & editing: S.V., M.J., S.L., S.G., M.O., M.J., K.J.A., A.G., C.J., F.P., J.M.C., S.B., M.T., A.P.S., A.S., M.R., S.H., and N.E. Funding acquisition: S.V., L.R.G., and N.E.

The authors declare that they have no conflict of interest.

## References

1. Max Roser, E. O.-O. & Ritchie, H. Life Expectancy. Our World Data (2020).

2. Partridge, L., Deelen, J. & Slagboom, P. E. Facing up to the global challenges of ageing. Nature 561, 45–56 (2018).

3. Dennison, E. M., Sayer, A. A. & Cooper, C. Epidemiology of sarcopenia and insight into possible therapeutic targets. Nat. Rev. Rheumatol. 13, 340–347 (2017).

4. Lappalainen, T. & Greally, J. M. Associating cellular epigenetic models with human phenotypes. Nat. Rev. Genet. 18, 441–451 (2017).

5. López-Otín, C., Blasco, M. A., Partridge, L., Serrano, M. & Kroemer, G. The Hallmarks of Aging. Cell 153, 1194–1217 (2017).

6. Horvath, S. & Raj, K. DNA methylation-based biomarkers and the epigenetic clock theory of ageing. Nat. Rev. Genet. 19, 371–384 (2018).

7. Zykovich, A. et al. Genome-wide DNA methylation changes with age in disease-free human skeletal muscle. Aging Cell 13, 360–366 (2014).

8. Day, K. et al. Differential DNA methylation with age displays both common and dynamic features across human tissues that are influenced by CpG landscape. Genome Biol. 14, R102 (2013).

9. Voisin, S. et al. An epigenetic clock for human skeletal muscle. J. Cachexia. Sarcopenia Muscle 11, 887–898 (2020).

10. Blocquiaux, S. et al. Recurrent training rejuvenates and enhances transcriptome and methylome responses in young and older human muscle. bioRxiv 2020.06.30.179465 (2020). doi:10.1101/2020.06.30.179465

11. Turner, D. et al. DNA methylation across the genome in aged human skeletal muscle tissue and muscle-derived cells: The role of HOX genes and physical activity. Sci. Rep. 10, 15360 (2020).

12. Willer, C. J., Li, Y. & Abecasis, G. R. METAL: fast and efficient meta-analysis of genomewide association scans. Bioinformatics 26, 2190–2191 (2010).

13. van Dongen, J. et al. Genetic and environmental influences interact with age and sex in shaping the human methylome. Nat. Commun. 7, 11115 (2016).

14. Dick, K. J. et al. DNA methylation and body-mass index: a genome-wide analysis. Lancet 6736, 1–9 (7AD).

15. Davegårdh, C., García-Calzón, S., Bacos, K. & Ling, C. DNA methylation in the pathogenesis of type 2 diabetes in humans. Mol. Metab. 14, 12–25 (2018).

16. Guo, S. et al. Identification of methylation haplotype blocks aids in deconvolution of heterogeneous tissue samples and tumor tissue-of-origin mapping from plasma DNA. Nat. Genet. 49, 635–642 (2017).

17. Vanderkraats, N. D., Hiken, J. F., Decker, K. F. & Edwards, J. R. Discovering high-resolution patterns of differential DNA methylation that correlate with gene expression changes. Nucleic Acids Res. 41, 6816–6827 (2013).

18. Schlosberg, C. E., VanderKraats, N. D. & Edwards, J. R. Modeling complex patterns of differential DNA methylation that associate with gene expression changes. Nucleic Acids Res. 45, 5100–5111 (2017).

19. Roadmap Epigenomics, C. et al. Integrative analysis of 111 reference human epigenomes. Nature 518, 317–330 (2015).

20. Zhou, W., Laird, P. W. & Shen, H. Comprehensive characterization, annotation and innovative use of Infinium DNA methylation BeadChip probes. Nucleic Acids Res. 45, e22–e22 (2017).

21. Fishilevich, S. et al. GeneHancer: genome-wide integration of enhancers and target genes in GeneCards. Database (Oxford). 2017, bax028 (2017).

22. Supek, F., Bošnjak, M., Škunca, N. & Šmuc, T. REVIGO Summarizes and Visualizes Long Lists of Gene Ontology Terms. PLoS One 6, e21800 (2011).

23. Su, J. et al. A novel atlas of gene expression in human skeletal muscle reveals molecular changes associated with aging. Skelet. Muscle 5, 35 (2015).

24. Ubaida-Mohien, C. et al. Discovery proteomics in aging human skeletal muscle finds change in spliceosome, immunity, proteostasis and mitochondria. Elife 8, (2019).

25. Greenberg, M. V. C. & Bourc’his, D. The diverse roles of DNA methylation in mammalian development and disease. Nat. Rev. Mol. Cell Biol. 20, 590–607 (2019).

26. Field, A. E. et al. DNA Methylation Clocks in Aging: Categories, Causes, and Consequences. Mol. Cell 71, 882–895 (2018).

27. Pérez, R. F., Tejedor, J. R., Bayón, G. F., Fernández, A. F. & Fraga, M. F. Distinct chromatin signatures of DNA hypomethylation in aging and cancer. Aging Cell 17, e12744 (2018).

28. Rakyan, V. K. et al. Human aging-associated DNA hypermethylation occurs preferentially at bivalent chromatin domains. Genome Res. 20, 434–439 (2010).

29. Jung, M. & Pfeifer, G. P. Aging and DNA methylation. BMC Biol. 13, 7 (2015).

30. Deschenes, M. R. Effects of aging on muscle fibre type and size. Sports Med. 34, 809–824 (2004).

31. Tchkonia, T. et al. Fat tissue, aging, and cellular senescence. Aging Cell 9, 667–684 (2010).

32. Cui, C.-Y. & Ferrucci, L. Macrophages in skeletal muscle aging. Aging (Albany. NY). 12, 3–4 (2020).

33. Brack, A. S. & Muñoz-Cánoves, P. The ins and outs of muscle stem cell aging. Skelet. Muscle 6, 1 (2016).

34. van Iterson, M., van Zwet, E. W. & Heijmans, B. T. Controlling bias and inflation in epigenome- and transcriptome-wide association studies using the empirical null distribution. Genome Biol. 18, 19 (2017).

35. Leek, J. T. et al. Tackling the widespread and critical impact of batch effects in high-throughput data. Nature reviews. Genetics 11, 733–739 (2010).

36. Pillon, N. J. et al. Transcriptomic profiling of skeletal muscle adaptations to exercise and inactivity. Nat. Commun. 11, 470 (2020).

37. Yan, X. et al. The Gene SMART study: Method, Study Design, and Preliminary Findings. BMC Genomics 18, 821 (2017).

38. Caiazzo, R. et al. Roux-en-Y gastric bypass versus adjustable gastric banding to reduce nonalcoholic fatty liver disease: a 5-year controlled longitudinal study. Ann. Surg. 260, 893–899 (2014).

39. Seaborne, R. A. et al. Human Skeletal Muscle Possesses an Epigenetic Memory of Hypertrophy. Sci. Rep. 8, 1898 (2018).

40. Ribel-Madsen, R. et al. Genome-wide analysis of DNA methylation differences in muscle and fat from monozygotic twins discordant for type 2 diabetes. PLoS One 7, e51302 (2012).

41. Gancheva, S. et al. Dynamic changes of muscle insulin sensitivity after metabolic surgery. Nat. Commun. 10, 4179 (2019).

42. Taylor, D. L. et al. Integrative analysis of gene expression, DNA methylation, physiological traits, and genetic variation in human skeletal muscle. Proc. Natl. Acad. Sci. 201814263 (2019). doi:10.1073/pnas.1814263116

43. Smyth, G. K. Limma: linear models for microarray data. (2005). doi:10.1007/0-387-29362-0_23

44. Peters, T. J. et al. De novo identification of differentially methylated regions in the human genome. Epigenetics {&} Chromatin 8, 1–16 (2015).

45. Kim, S., Yu, N.-K. & Kaang, B.-K. CTCF as a multifunctional protein in genome regulation and gene expression. Exp. Mol. Med. 47, e166–e166 (2015).

46. Margueron, R. & Reinberg, D. The Polycomb complex PRC2 and its mark in life. Nature 469, 343–349 (2011).

47. Phipson, B., Maksimovic, J. & Oshlack, A. missMethyl: an R package for analyzing data from Illumina’s HumanMethylation450 platform. Bioinformatics 32, 286–288 (2016).

48. Maksimovic, J., Oshlack, A. & Phipson, B. Gene set enrichment analysis for genome-wide DNA methylation data. bioRxiv 2020.08.24.265702 (2020). doi:10.1101/2020.08.24.265702

49. Benjamini, Y. & Hochberg, Y. Controlling the false discovery rate: a practical and powerful approach to multiple testing. J Roy Stat Soc B Met 57, 289–300 (1995).

50. Benjamin, D. J. et al. Redefine Statistical Significance. Hum. Nat. Behav. 2, 6–10 (2017).

